# Kinetic characteristics of propofol-induced inhibition of electron-transfer chain and fatty acid oxidation in human and rodent skeletal and cardiac muscles

**DOI:** 10.1101/633156

**Authors:** Tomáš Urban, Petr Waldauf, Adéla Krajčová, Kateřina Jiroutková, Milada Halačová, Valér Džupa, Libor Janoušek, Eva Pokorná, František Duška

## Abstract

**Introduction:** Propofol causes a profound inhibition of fatty acid oxidation (FAO) and reduces spare electron transfer chain (ETC) capacity in a range of human and rodent cells and tissues – a feature that might be related to the pathogenesis of Propofol Infusion Syndrome. We aimed to explore the mechanism of propofol-induced alteration of bioenergetic pathways by describing its kinetic characteristics.

**Methods:** We obtained samples of skeletal and cardiac muscle from Wistar rat (n=3) and human subjects: *vastus lateralis* from hip surgery patients (n=11) and myocardium from brain-dead organ donors (n=10). We assessed mitochondrial functional indices using standard SUIT protocol and high resolution respirometry in fresh tissue homogenates with or without short-term exposure to a range of propofol concentration (2.5-100 μg/ml). After finding concentrations of propofol causing partial inhibition of a particular pathways, we used that concentration to construct kinetic curves by plotting oxygen flux against substrate concentration during its stepwise titration in the presence or absence of propofol. By spectrophotometry we also measured the influence of the same propofol concentrations on the activity of isolated respiratory complexes.

**Results:** We found that human muscle and cardiac tissues are more sensitive to propofol-mediated inhibition of bioenergetic pathways than rats tissue. In human homogenates, palmitoyl carnitine-driven respiration was inhibited at much lower concentrations of propofol than that required for a reduction of ETC capacity, suggesting FAO inhibition mechanism different from downstream limitation or carnitine-palmitoyl transferase-1 inhibition. Inhibition of Complex I was characterised by more marked reduction of Vmax, in keeping with non-competitive nature of the inhibition and the pattern was similar to the inhibition of Complex II or ETC capacity. There was no inhibition of Complex IV nor increased leak through inner mitochondrial membrane with up to 100 μg/ml of propofol. If measured in isolation by spectrophotometry, propofol 10 μg/ml did not affect the activity of any respiratory complexes.

**Conclusion:** In human skeletal and heart muscle homogenates, propofol in concentrations that are achieved in propofol-anaesthetized patients, causes a direct inhibition of fatty acid oxidation, in addition to inhibiting flux of electrons through inner mitochondrial membrane. The inhibition is more marked in human as compared to rodent tissues.

## Introduction

Propofol is a short-acting hypnotic agent, which is reportedly administered to 100 millions of patients each year and which was has been on the WHO list of most essential drugs for more than 10 years[1]. However, in last three decades, fatal complications of propofol administration were reported and defined as Propofol infusion syndrome (PRIS)[2,3]. The syndrome typically includes metabolic acidosis, arrhythmias, ECG changes, hyperlipidaemia, fever, hepatomegaly, rhabdomyolysis, cardiac and/or renal failure. Risk of developing PRIS increases with higher dose and duration of infusion[4], and several studies in animals[5–9] and humans[10], [11–13] suggest that this syndrome might be an extreme manifestation in susceptible individuals of propofol inhibitory effect on fatty acid oxidation (FAO) and/or electron-transfer chain. Vanlander et al. in a study on rodents[14] first brought evidence for a hypothesis, that due to structural similarity of propofol and Coenzyme Q (CoQ), at least some effects of propofol on bioenergetics are caused by the inhibition of CoQ-dependent electron transfer pathways. Other mechanisms are possible, too, including metabolic rearrangement at translational level [15,16]. It is unknown whether increased concentration of substrates can overcome propofol-induced inhibition, a feature that would point towards competition of propofol with the respective substrate, or whether the presence of propofol influences the maximum flux through the ETC, which would point towards inhibition outside the substrate-binding sites of the enzymes or an interruption of electron flux through the respiratory chain, such as at the level of CoQ.

The insight into the mechanisms of propofol toxicity was mostly obtained from experiments in animals and it is currently unknown whether effects of propofol can be reproduced in human tissues. In this study, we assessed propofol-effect on energy metabolism in skeletal and heart muscle in humans and rats. We aimed to (1.) clarify the inter-species differences in the effect of propofol and (2.) elucidate the Michaelis-Menten characteristics of propofol-induced inhibition of metabolic pathways that will have been found to be inhibited by propofol. In order to maximise biological plausibility of our results, we performed our key experiments on tissue homogenates containing mitochondrial networks in cytosolic context[17].

## Methods

### Skeletal and heart muscle samples

Overall study design is shown in Fig 1. All experiments were performed ex vivo, with all the respirometry measurement being done on fresh tissue homogenates. Human skeletal muscle biopsies were obtained from *m. vastus lateralis* (300 mg) from metabolically healthy volunteers (n=11, 4 males, 7 females, aged 63±10 years) undergoing hip replacement surgery at Department of Orthopaedic Surgery in FNKV University Hospital in Prague. We excluded both patients receiving propofol anaesthesia during surgery and patients with history of mitochondrial disorder, diabetes mellitus or any other metabolic disease except treated stable hypothyroidism. Human heart muscle biopsies (100 mg) were taken from left ventricle myocardium from brain-dead donors (n=10) free of cardiac disease, whose hearts were not suitable for transplantation due to donor age over 50 years or due to logistical reasons, at the Transplant Centre of Institute of Clinical and Experimental Medicine in Prague, Czech Republic. All skeletal muscle donors gave a prospective written informed consent. In brain dead donors, organ retrieval consent process was performed in accordance with Czech legal and ethical requirements and the need for specific informed consent for this study was waived by Research Ethics Boards in both participating institutions, which also reviewed and approved the protocol of the study. Detailed characteristics of brain dead donor subjects are in Supplementary appendix Table S1.

**Figure 1.**
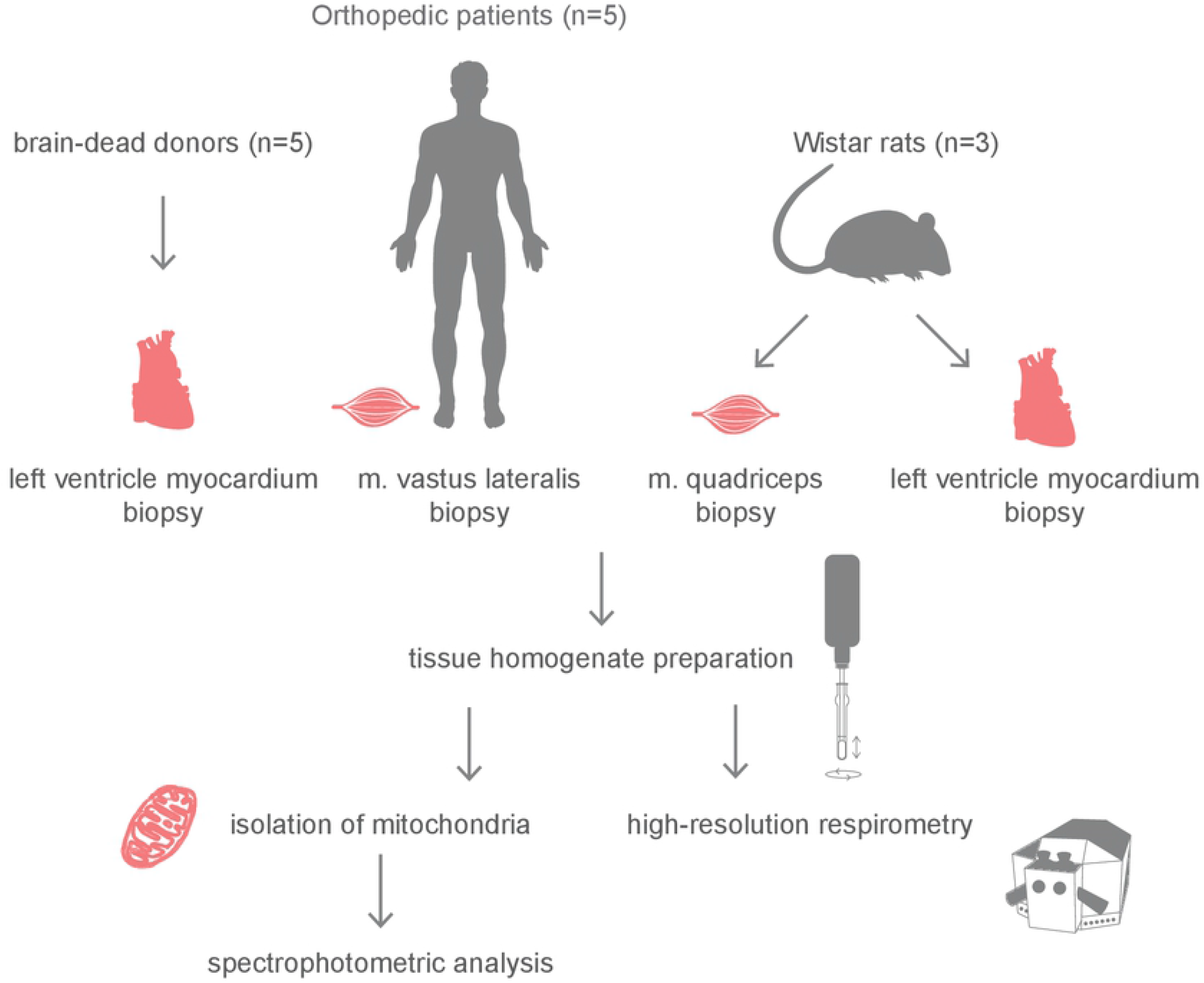
Study protocol overview.

Rat muscle tissue samples (100-200 mg) were obtained from male Wistar rats (n=3). The rats were anesthetized via inhalation of diethylether and sacrificed by decapitation. Tissue samples were retrieved immediately before homogenisation from skeletal muscle *m. quadriceps femoris* and left ventricle myocardium. The design of this part of the protocol was reviewed and approved by the Committee for Protection and Care of Animals Used in Medical Research of the National Institute of Public Health.

### Preparation of tissue homogenates

Muscle biopsy samples were put in pre-cooled biopsy preservation and relaxing medium (BIOPS) consisting of 2.77 mM CaK_2_EGTA, 7.23 mM K_2_EGTA, 5.77 mM Na_2_ATP, 6.56 mM MgCl_2_ x 6 H_2_O, 20 mM taurine, 15 mM Na_2_Phosphocreatine, 20 mM imidazole, 0.5 mM dithiothreitol and 50 mM MES hydrate (pH adjusted to 7.1 at 4°C)[18]. Skeletal and heart muscle homogenates were prepared according to the protocols previously described by our research group^14,17^ and in the step-by-step protocol in Supplementary Appendix. Briefly, in skeletal muscle samples, the connective and adipose tissue was removed using scissors and forceps under microscope and muscle sample was gently blotted by sterile cotton gauze, weighted on the analytical scale and cut into fine fragments. Tissue fragments were then placed into the glass grinder of 2 mL Potter-Elvehjam homogenizer set (Wheaton™, Millville, USA) and diluted in the respiration medium (K medium) containing 80 mM KCl, 10 mM Tris HCl, 5mM KH_2_PO_4_, 3 mM MgCL_2_, 1 mM EDTA, 0.5 mg/mL BSA (pH adjusted to 7.4 at 24 °C) to obtain 10 and 2.5 % tissue solution (100 mg of human and 25 mg of rat skeletal muscle tissue /per 1 mL of K medium, respectively). Tissue fragments were then homogenized by 5-6 slow strokes up and down with 2 mL Potter-Elvehjam teflon/glass homogenizer (clearance 0.25 mm; Wheaton™, Millville, USA) driven by electric motor homogenizer (750 rpmi; HEi-Torque Value 100, Heidolph, Germany)[17]. Heart muscle homogenates were prepared similarly[19]. After removal of connective tissue and fat, fresh myocardium was dissected into fine fragments and 2.5 and 1% tissue solution (25 mg of human and 10 mg of rat heart muscle tissue /per 1 mL) was obtained when diluting in a respiration medium of different composition (MiR05): 0.5 mM EGTA, 3 mM MgCl_2_ x 6 H_2_O, 60 mM lactobionic acid, 20 mM taurine, 10 mM KH_2_PO_4_, 20 mM HEPES, 110 mM sucrose and 1 g/L of BSA (pH adjusted to 7.1 at 24°C). Then the process of homogenization was two-step: tissue fragments were firstly manually homogenized by 10-12 initial strokes up and down with a glass loose pestle (large clearance 0.114 ± 0.025 mm; Wheaton™, Millville, USA) in the glass Dounce grinder and subsequently homogenized with 5-6 slow strokes by a motor-driven PTFE pestle of 2 mL Potter-Elvehjam homogenizer (750 rpmi; Wheaton™, Millville, USA)[19]. Finally, crude homogenate was filtered through polyamide mesh (SILK & PROGRESS s.r.o., Czech Republic) in order to remove the remaining connective tissue. All the steps were performed on ice. Integrity of outer mitochondrial membrane was verified by measuring the change of oxygen flux at the presence of ADP after addition of Cytochrome c. In all experiments, the increments were <15%.

### High resolution respirometry

All functional studies were performed using high resolution respirometry Oxygraph-2k (O2k; Oroboros Instruments, Innsbruck, Austria). The method is based on monitoring oxygen concentration with a polarographic oxygen electrode in two closed chambers allowing for 2 parallel measurements. Oxygen flux is calculated as the negative derivative of oxygen concentration[20,21] which is continually measured during sequential addition of substrates, uncouplers or inhibitors. In our experiments, we performed all initial steps including washing the chamber and calibration of O2k according to the manufacturer’s recommendations[22]. Oxygen solubility factor was set up to 0.93 and 0.92 for K medium and MiR05, respectively. Changes of respiration were recorded and analysed by Datlab software (Datlab Version 7.0, Oroboros Instruments, Innsbruck, Austria). See Supplementary appendix for detailed methodology.

### Reagents preparation

All reagents were purchased from Sigma-Aldrich (St. Louis, USA). Substrates were dissolved in distilled water (pH adjusted to 7.0). Uncouplers and inhibitors were dissolved in DMSO. Propofol (2,6-diisopropyplhenol) stock (10 mg/mL) was prepared fresh before each measurement by dilution in 10 % ethanol.

### Substrate-uncoupler-inhibitor-titration (SUIT) protocols for dose-finding studies

In the dose-finding studies, we tested a range of propofol concentrations, from those resembling propofol levels in human plasma during anaesthesia and sedation (2.5; 5 and 10 μg/mL)[23,24] to supra-physiological concentrations used in previous animal studies (25; 50 and 100 μg/mL)[5–7,25]. Propofol or the respective concentration of control vehicle (ethanol) were injected directly into the chamber and both measurements were performed simultaneously.

#### Global mitochondrial functional indices

In the first two experiments, we assessed propofol influence on global bioenergetic parameters. *Inhibition of electron transfer chain (ETC)*. Firstly, we used a sequential addition of 2.5 mM malate (mal) plus 15 mM glutamate (glut), 1 mM ADP, 10 μM cytochrome c, 10 mM succinate (suc), 1 μM oligomycin, 0.8 and 1.5 μM FCCP (for skeletal muscle and heart muscle homogenates, resp.) followed by final injection of increasing concentrations of either propofol or vehicle. This protocol allowed us to assess possible propofol-induced inhibition of ETC in ADP-stimulated respiration. *Uncoupling of inner mitochondrial membrane (Leak)*. Secondly, we looked at possible uncoupling of inner mitochondrial membrane and changed the protocol as follows: addition of 2.5 mM malate plus 15 mM glutamate, 1 mM ADP, 10 μM cytochrome c, 10 mM succinate and 1 μM oligomycin was followed by final titration of propofol concentrations to induce feasible uncoupling (Fig. 2).

**Figure 2.**
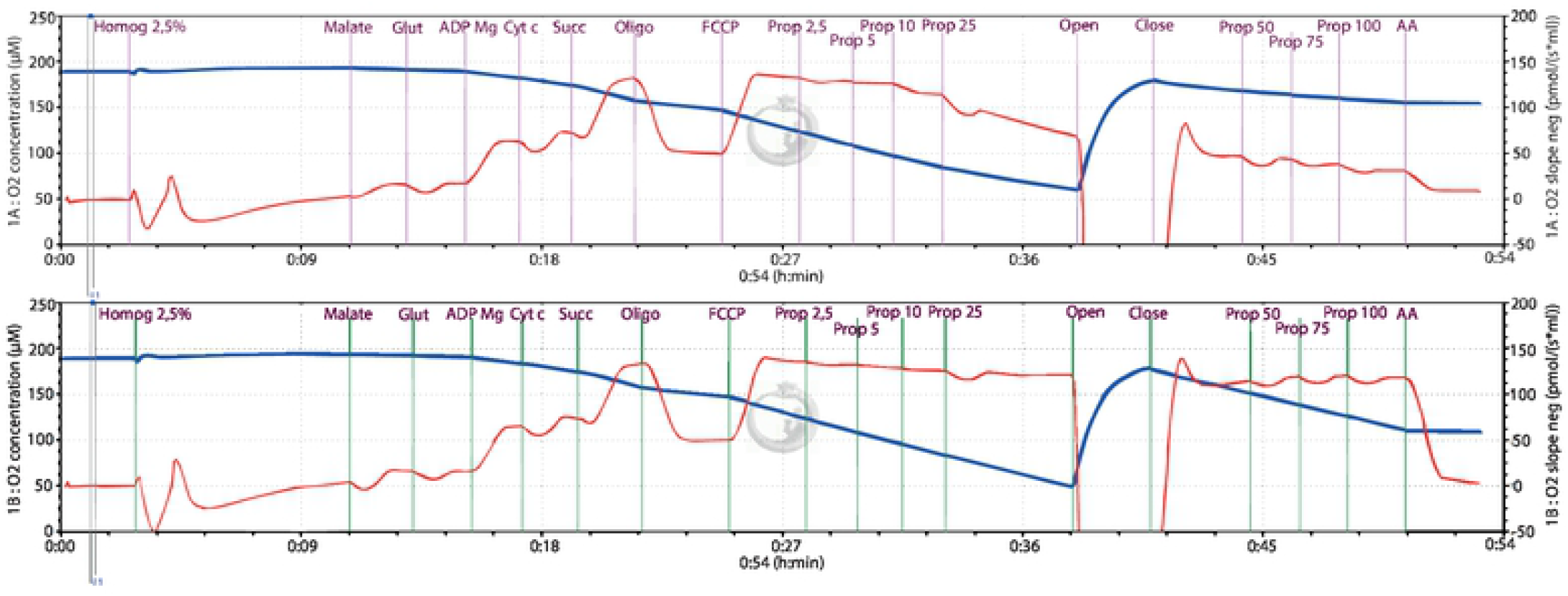
Example of SUIT high resolution respirometry protocol. The two measurements performed in the presence of propofol (μg/ml) or vehicle (ethanol) as control.

#### Respiration linked to individual complexes

In experiments 3-5 we focused on propofol-induced changes in complexes I, II and IV of the ETC that were assessed by high resolution respirometry in muscle tissue homogenates. *Complex I-linked respiration*. Firstly, 5 mM malonate, a specific inhibitor of complex II, and 2.5 mM malate plus 15 mM glutamate as substrates for complex I were injected followed by addition of 1 mM ADP and 10 μM cytochrome c. Lastly, propofol was added in a step-wise manner. *Complex II-linked respiration*. Firstly, 3.5 μM rotenone was added to block complex I-linked respiration and 10 mM succinate was added as substrate for complex II followed by 1 mM ADP and 10 μM cytochrome c. Again, the experiment was finished by step-wise addition of propofol *Complex IV-linked respiration*. Upstream electron flux was blocked by 4 μM antimycin A, a specific inhibitor of complex III. Then, artificial Complex IV substrate 10 mM ascorbate plus 0.2 mM TMPD together with 1 mM ADP and 10 μM cytochrome c were added to maximise complex IV-linked respiration followed by stepwise addition of increasing concentrations of propofol.

#### Fatty acid oxidation (FAO)

Initially, 2.5 mM malate was used as a sparkler followed by 1 mM ADP. Respiration was then stimulated by addition of 25 μM of palmitoyl-carnitine (heated to 70°C). Lastly, propofol was added in a step-wise manned to observe its potential influence on FAO.

### Kinetic characteristics experiments

Because we found propofol to inhibit CI-driven respiration and FAO in concentrations that are clinically relevant, for these pathways we performed substrate titrations in parallel in control chamber and in the presence of a fixed concentration of propofol (10 μg/ml in rodents and 2.5 and 10 μg/ml in humans). For Complex I-linked respiration we inhibited complex II by malonate, added ADP, cytochrome c, and propofol after which we started to add malate/glutamate (CI substrates) in a stepwise manner (0.025/0.15; 0.1/0.3; 0.25/1.5; 1/3; 2.5/15 mM). In analogy, to study propofol-induced FAO inhibition we first added malate and ADP, then propofol and then titrated palmitoyl-carnitine (0.1; 0.25; 1; 2.5; 10; 25 uM). Oxygen flux was expressed as % of respiration in the control chamber and plotted against substrate concentration.

### Spectrophotometric analyses of isolated respiratory complexes

In order to assess effects of propofol on isolated complexes I, II, III and IV, we used classical spectrophotometric method on frozen homogenates as described by Spinazzi et al[26]. We performed parallel assays of samples containing 10 μg/ml of propofol, the solvent (ethanol 0.5 %) and fresh media. For measuring of *complex I activity* we used solution of 0.05 M potassium phosphate buffer, 3 mg/ml BSA, 0.3 mM KCN, 0.1 mM NADH, added isolated mitochondria and measured it in 96 well plate. We used parallel measurement with and without addition of 10μM rotenone, read the baseline at 340 nm for 2 minutes and started reaction by injection ubiquinone, with absorbance 340 nm duration time of 2 minutes. For *Complex II activity* measurement we used solution of 0.05 M potassium phosphate buffer, 1 mg/ml BSA, 0.3 mM KCN, 20 mM succinate and added isolated mitochondria. Baseline was read at 600 nm for 3 minutes then we started the reaction with 12.5 mM decylubiquinone, checking specificity with 1 M malonate. For measurement *complex III* activity we used solution of 0.05 M potassium phosphate buffer, 75 μM oxidized cytochrome c, 0.1 mM EDTA, 2.5% Tween-20 and isolated mitochondria. Read baseline at 550 nm for 2 minutes and start the reaction with 10 mM decylubiquinone and check specificity with 10 mM KCN. *Activity of complex IV* was measured with solution of 0.05 M potassium phosphate buffer, 60 μM reduced cytochrome c then the baseline was read at 550 nm for 2 minutes. Reaction was start by adding isolated mitochondria and the specificity was checked by adding 0.3 mM KCN in one of the measurements.

### Statistics

Data from dose-finding were processed using linear mixed effect model (LMEM). In the fixed part, the model consists of a dependent continuous parameter (e.g. oxygen flux) and a categorical independent parameter (e.g. propofol concentration, species human vs rodent). In the random part, there was a random intercept (ID of a patient) and correlation matrix of residuals, if statistically significant (as assessed by likelihood ratio test). For kinetic characterization of propofol inhibition of complex I-linked respiration and FAO, we used mixed effect multilevel non-linear regression for finding the best fit into Michaelis-Menten equation, from which we calculated Km and Vmax. For all statistical analyses we used software Stata 15.1 (Stata Corp., LLC, U.S.A.).

## Results

### Effect of propofol on bioenergetics in cytosolic context

Propofol inhibited in a concentration-dependent manner oxygen fluxes driven by substrates for CI and by palmitoyl-carnitine, and to lesser extent of oxygen fluxes driven by substrate for Complex II (Fig. 3). In turn, respiratory chain capacity (RCC) was reduced by propofol, too. Skeletal muscle (both of man and rat) seemed to be more sensitive to inhibition of CI-driven respiration by propofol than myocardium. In contrast, CII-driven respiration, FAO and RCC were first inhibited in human myocardium during propofol titration. The inhibition of FAO and CI-driven respiration were inhibited at concentrations of propofol seen in plasma of anaesthetized patients, i.e. 2.5 and 10 μg/ml and human tissues seemed to be more sensitive than rodents’. Apparently, there was no significant effect of propofol on Complex IV-driven respiration or on the leak of electrons through inner mitochondrial membrane.

**Figure 3.**
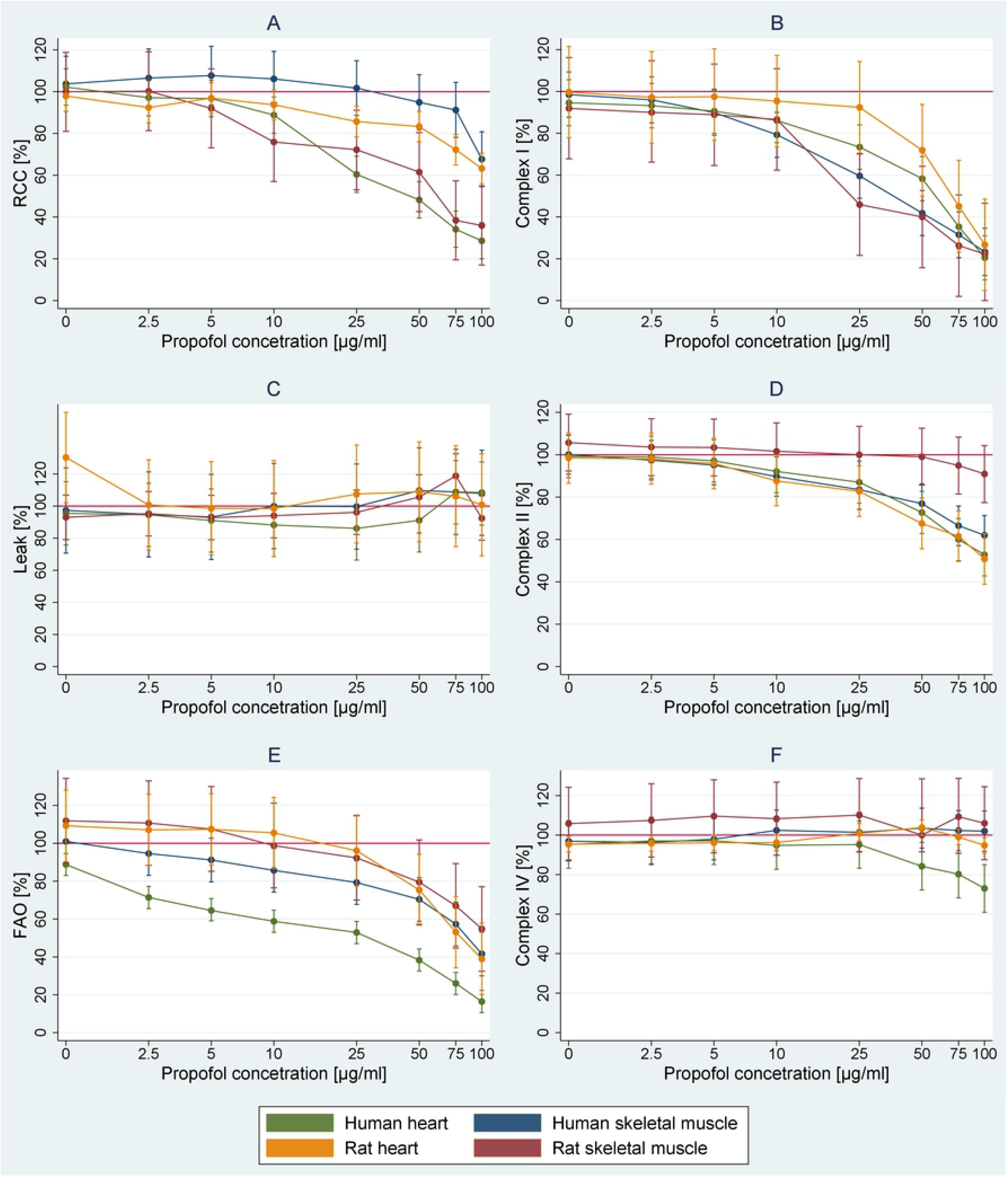
Prediction of difference of respiratory parameters against control at different concentration of propofol. Graph A – respiration chain capacity, graph B – complex I, graph C – LEAK, graph D - complex II, graph E – FAO, graph F – complex IV.

In light of this we selected CI-driven and palmitoyl carnitine-driven respirations for kinetic studies, in which we use 10 μg/ml (for human tissues 2.5 and 10 μg/ml, respectively) of propofol.

### Kinetic characteristics of inhibition by propofol

Energy substrates were titrated with or without the presence of propofol and pre- and post-inhibition Vmax and Km were compared. (See Figures 2 and 4). Propofol influenced kinetic parameters of Complex I-driven respiration in human, but not rat muscle tissues: it reduced Vmax and to a lesser, but still significant extend, the Km. At 2.5 μg/ml, only Vmax in human skeletal muscle was significantly affected (Fig. 4A,B). With regards to FAO, even the lower concentration (2.5 μg/ml) of propofol significantly reduced both Km and Vmax in both human skeletal muscle and heart tissues. The higher concentration (10 μg/ml) also tended to reduce Vmax in rat tissues, but the reduction was only significant in skeletal muscle (Fig. 4 C,D).

**Figure 4.**
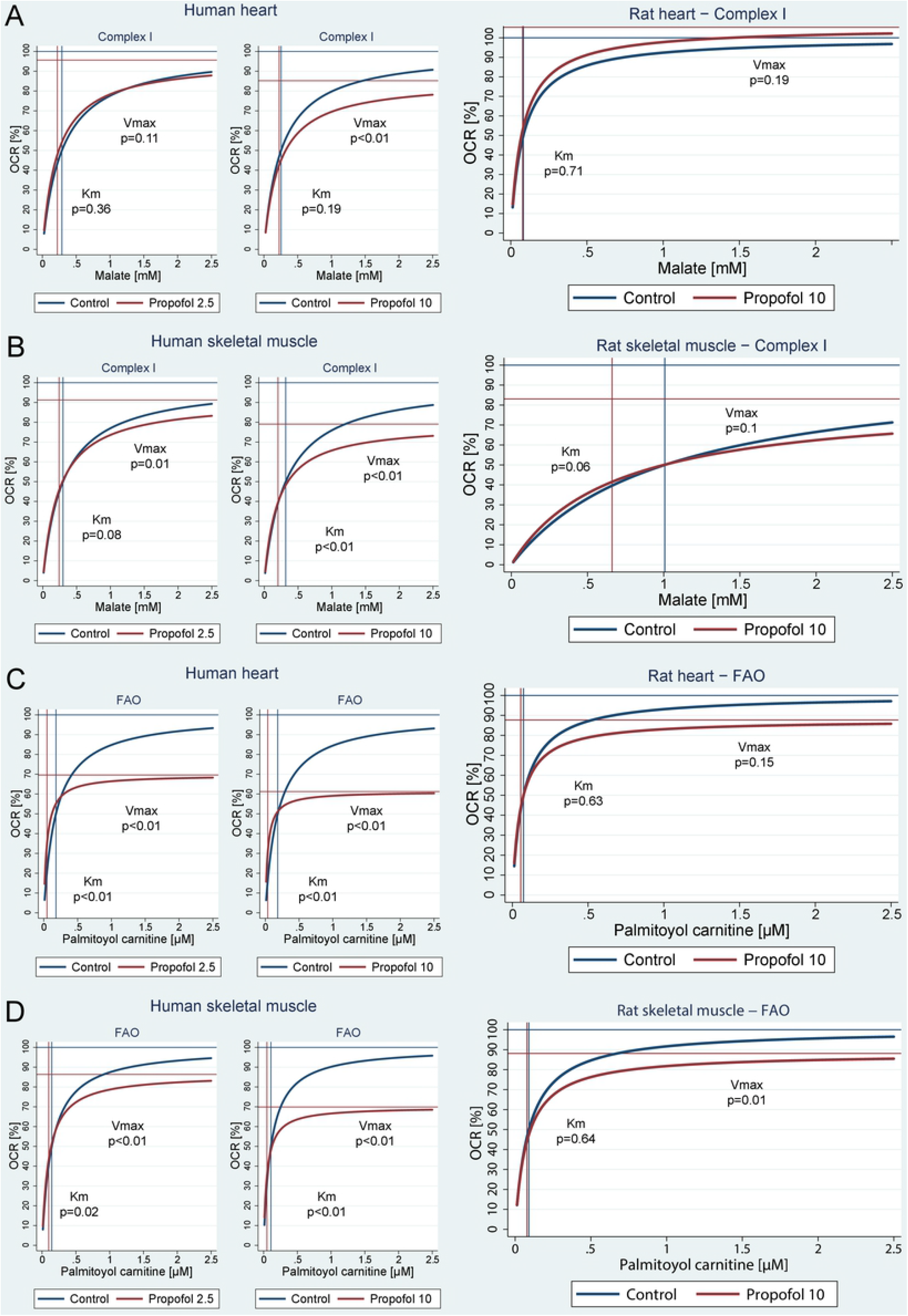
Kinetics of inhibition by 2.5 μg/ml and 10 μg/ml of propofol in human muscle homogenates (left and middle figures), and by 10 μg/ml in rat muscle homogenates (right figure). A = Complex I in heart tissue. B = Complex I in skeletal muscle. C= FAO in heart tissue. D = FAO in skeletal muscle. Based on multi-level linear regression model of independent measurements in all subjects. Note: FAO = fatty acid oxidation, OCR = oxygen consumption rate.

### Effects of propofol on isolated complexes

Of note there was no significant inhibition by 10 μg/ml of propofol of any of the complexes if measured in isolation by spectrophotometry (See Supplementary Appendix, Fig. S1).

## Discussion

To our knowledge this is the first study that investigates the effects of clinically relevant concentration of propofol on cellular bioenergetics in human skeletal and heart muscles obtained from volunteers and brain-dead organ donors. The main finding of our study is that human skeletal and cardiac muscle tissues are more sensitive to bioenergetic effects of propofol when directly compared to rodent samples. Palmitoyl-carnitine driven respiration (FAO) is particularly more sensitive to low concentrations of propofol in humans compared to rat (Fig. 3E). In addition, we have found that in the model of short-term exposure to propofol, there is a different pattern of inhibition in Complex I-driven respiration from palmitoyl carnitine-driven respiration. In our experiments, we measured the rate of oxygen consumption during substrate titration in the presence or absence of propofol. If propofol competed at the binding site of any substrate or intermediate, thus creating a bottleneck in the reaction, increasing the concentration of substrate would have led to the same maximal reaction velocity (Vmax), but indeed the concentration at which 50% Vmax is achieved (i.e. Km) would have been higher. On the contrary, if propofol inhibited the rate-limiting enzyme a non-competitive way (e.g. by binding outside the binding site or draining electrons from the ETC as proposed[14]), Km would remained unchanged and only Vmax would decrease. The patterns we have observed for complex-I-driven respiration and palmitoyl carnitine-driven respiration (FAO) are different. Complex-I-driven respiration in human skeletal muscle is inhibited by much lower propofol concentrations compared to those that are able to influence overall capacity of ETC (see Fig 3A, 3B). Km long remains unaffected and only a reduction of Vmax is seen (see Fig. 4B), in keeping with a non-competitive nature of the inhibition. Because the activity of isolated complexes was unaffected by the same concentration of propofol (10 μg/ml), inhibition of Complex I-driven respiration could be explained by propofol reducing electron flux outside binding sites of the enzymes, such as reducing downstream flow of electrons by interfering with Coenzyme Q. On the contrary, palmitoyl carnitine-driven respiration was affected differently by propofol: both Vmax and Km were reduced (Fig. 4C and 4D) suggesting a pattern of mixed or uncompetitive inhibition. Because FAO was inhibited even by concentrations <5 μg/ml (Fig. 3E) which are not high enough to cause any measurable reduction of ETC capacity (Fig 3A) or Complex II substrate-driven respiration (Fig. 3D), the reduction of downstream flux of electrons by propofol can be effectively ruled out as the only cause of FAO inhibition. Propofol has long been thought to inhibit carnitine-palmitoyl transferase I (CPT-1) after Wolf[11] reported in the blood of a child with PRIS a significant elevation of acyl-CoA including malonyl-CoA, the main physiological inhibitor of CAP-1 and in turn the main regulator of fatty acids transport into the mitochondrial matrix. Of note, CAP-1 converts acyl-CoA to acyl-carnitine and in this experiment, we used palmitoyl-carnitine as the substrate. In turn, the observed inhibition of FAO cannot be explained by effects of propofol on CPT-1 as often reported[27,28]. According to our data, the inhibition of FAO by propofol in human skeletal and cardiac muscle cell must occur between CPT-2 and entry of electrons from FAO into ETC via Complex II, and not at the level of CPT-1.

Our results complement the finding of Branca et. al.[5] and Rigoulet et al.[25] who found inhibition of ETC capacity in isolated rat liver, mostly due to inhibition of Complex I. In keeping with our results, rat heart mitochondria were found to be more resistant to the inhibitory effects of propofol as the effects of propofol were only seen with concentrations > 55 μg/ml (>300 μmol/l)[6]. Rare in vivo studies in guinea pigs[29] or rat [14] demonstrated – in line with our results – the inhibitory effect of propofol in ranges of 9-36 μg/ml (50-200 μmol/l)[29] or >4.5 μg/ml (>25 μmol/l)[14], respectively, whilst isolated respiratory complexes were also unaffected by the same concentration of propofol[14]. Vanlender et al. was also able to mitigate propofol toxicity by adding Coenzyme Q to tissue homogenate[14] – a result we were unable to reproduce neither with Coenzyme Q nor with its analogue idebenone, mainly due to the toxicity of its solvent dimethylsulfoxide (data not shown). Yet, the kinetic characteristics of inhibition of Complex I-driven respiration in human tissues observed in this study, is in keeping with Coenzyme Q hypothesis.

The major strength of our work is that we directly measured the effects of propofol on human tissues that are clinically affected by PRIS, such as skeletal muscle (rhabdomyolysis) or myocardium (heart failure, arrhythmias). Native human hearts are very difficult to obtain and we used hearts retrieved from brain-dead organ donors to preform our experiments. We experimented with concentrations of propofol that are found in blood of patients sedated or anaesthetized with propofol (2-11 μg/ml)[24] and used biologically plausible bioenergetic model of mitochondria in their cytosolic context[17]. The finding of the differences between human and rodent tissues exposed to propofol we consider very important for the interpretation of the experiments in animal models of PRIS[5,7,14,26,29,30]. On the other hand, the use of muscle homogenates only allows short term experimental exposure[17], which may not allow sufficient time for incorporation of propofol into the inner mitochondrial membrane[14] or affect gene expression[15,16]. Also, measuring parameters from the classic enzyme kinetics (such as Vmax or Km) may not be fully appropriate in systems that are likely multicompartmental and complex, such as bioenergetic pathways in mitochondria.

In conclusion, we demonstrated for the first time in fresh human skeletal muscle and cardiac tissue homogenates the inhibition by propofol of electron transfer chain and fatty acid oxidation. We have shown that human tissues are more sensitive to propofol than rat and that fatty acid oxidation in humans is very sensitive to an inhibition by propofol and occurs at much lower concentrations, before any observable reduction of electron flux through ETC. Kinetic characteristics of fatty acid oxidation suggest a different mechanism of inhibition from previously proposed CTP-1 inhibition. Noncompetitive nature of the inhibition of Complex I substrate-driven respiration is consistent with the hypothesis that propofol effect on mitochondria are mediated by its interference with Coenzyme Q.

## Acknowledgement

Study was supported by grants AZV 16-28663A and Institutional Support for FNKV University Hospital from Czech Ministry of Health and by PROGRES Q37. We thank to all the volunteers who agreed to participate in this study and grieved families of organ donors for their gift of life. Dr Zdeněk Drahota provided invaluable advice and Přemysl Kunčický and Jiří Bukovský helped to carry out pilot experiments.

